# No kissing in the nucleus: Unbiased analysis reveals no evidence of trans chromosomal regulation of mammalian immune development

**DOI:** 10.1101/212985

**Authors:** Timothy M. Johanson, Hannah D. Coughlan, Aaron T.L. Lun, Naiara G. Bediaga, Gaetano Naselli, Alexandra L. Garnham, Leonard C. Harrison, Gordon K. Smyth, Rhys S. Allan

## Abstract

It has been proposed that interactions between mammalian chromosomes, or transchromosomal interactions (also known as kissing chromosomes), regulate gene expression and cell fate determination. Here we aimed to identify novel transchromosomal interactions in immune cells by high-resolution genome-wide chromosome conformation capture. Although we readily identified stable interactions in *cis,* and also between centromeres and telomeres on different chromosomes, surprisingly we identified no gene regulatory transchromosomal interactions in either mouse or human cells, including previously described interactions. We suggest that advances in the chromosome conformation capture technique and the unbiased nature of this approach allow more reliable capture of interactions between chromosomes than previous methods. Overall our findings suggest that stable transchromosomal interactions that regulate gene expression are not present in mammalian immune cells and that lineage identity is governed by *cis,* not *trans* chromosomal interactions.

## Introduction

Each chromosome contains just one DNA molecule. Recent technological advances have allowed characterisation of the elaborate three-dimensional structures that form from this DNA (*1).* These structures include globules or domains, which partition the chromosome, and elegant DNA loops that link gene promoters to distant enhancers. In addition to these intrachromosomal structures formed within the same DNA molecule, there are transchromosomal interactions formed between different chromosomes. Relative to intrachromosomal interactions, the frequency, nature and function of transchromosomal interactions are poorly understood (2).

In contrast to the multitude of intrachromosomal interactions known to regulate gene expression, only a handful of transchromosomal interactions have been described. For example, transchromosomal interactions were reported to be crucial for the appropriate expression of a single olfactory gene amongst the ~1300 within the genome (*3, 4*) and for X chromosome inactivation (*5–7).*Interestingly, a large number of the reported transchromosomal interactions have been characterised in cells of the immune system. For example, in both mouse and human T cells the insulin like growth factor 2 (Igf2) locus was reported to interact with a number of loci on different chromosomes (*8–10).* Also in T cells, a regulatory region on mouse chromosome 11 (the T helper 2 locus control region; LCR) was suggested to interact with loci encoding the cytokine interferon gamma (Ifng) on chromosome 10 (11) and interleukin 17 (IL-17) on chromosome 1 (12). Perturbation of these interactions was associated with altered expression of Ifng and IL-17, respectively. In mouse B cell progenitors, the interaction between the immunoglobulin heavy chain (*Igh*) locus on chromosome 12 and the immunoglobulin light chain (*Igk*) locus on chromosome 6 was important for the rearrangement of the heavy chain locus (13).

These transchromosomal interactions were all identified by either chromatin conformation capture, in which crosslinking, dilution of a ligation reaction and PCR are used to deduce the relative physical proximity of two loci in three-dimensions, or DNA FISH in which microscopy and labelled probes are used to locate loci within individual nuclei, or both. These techniques are targeted approaches. Here we aimed to use an unbiased, genome-wide approach to identify novel gene regulatory transchromosomal interactions in three distinct mouse and human immune cell populations. Unexpectedly, we found very few interactions between chromosomes, and none were gene regulatory or conserved. Overall, our findings question the existence of stable, gene-regulatory transchromosomal interactions underlying immune cell identity.

## Results

To elucidate novel transchromosomal interactions, we generated *in situ* HiC libraries from both mouse and human B cells and CD4^+^ and CD8^+^ T cells of the immune system (Supplemental Figure 1 A, B). The resulting ~200 million paired-end reads were then mapped to the appropriate genome, filtered for artefacts, such as dangling ends and self-circling reads, and counted into 50kb bins with the *diffHic* software package (14). After iterative correction of the generated contact matrices, DNA-DNA interactions were detected by comparing the interaction intensity in each bin to those surrounding it to determine significant interactions relative to background (15).

Using this pipeline we detected hundreds of interactions between chromosomes in each cell population (Supplemental Table 1). Consistent with previous literature (16), these transchromosomal interactions are enriched in gene-rich, centrally located chromosomes (Figure 1 A, Supplemental Figure 1 C). However, closer examination of these interactions reveals that a high percentage (74-90% in mouse and 82-94% in human) contain regions recommended to be removed, or ‘blacklisted׳, from analyses due to their high or low mappability, repeated nature, location within telomeres or centromeres, among others (*17, 18).* After application of blacklisting the majority of transchromosomal interactions are removed (Figure 1 B-C, Supplemental Table 2). This is in stark contrast to intrachromosomal interactions, of which less than 3% contain blacklisted regions (Figure 1 D). The majority of transchromosomal interactions remaining after blacklisting linked regions close to telomeres (Figure 1 E, F, Supplemental Figure 1 D, E) or centromeres (Figure 1 E, G, Supplemental Figure 1 F, G). Telomeres and centromeres physically cluster not just during preparation for mitosis, but throughout the cell cycle (19). Thus it appears that the majority of the transchromosomal interactions detected in mammalian immune cells may be a consequence of this telomeric and centromeric clustering. Importantly, the detection of these interactions confirms that *in situ* HiC is able to detect interactions between chromosomes.

**Figure 1:**
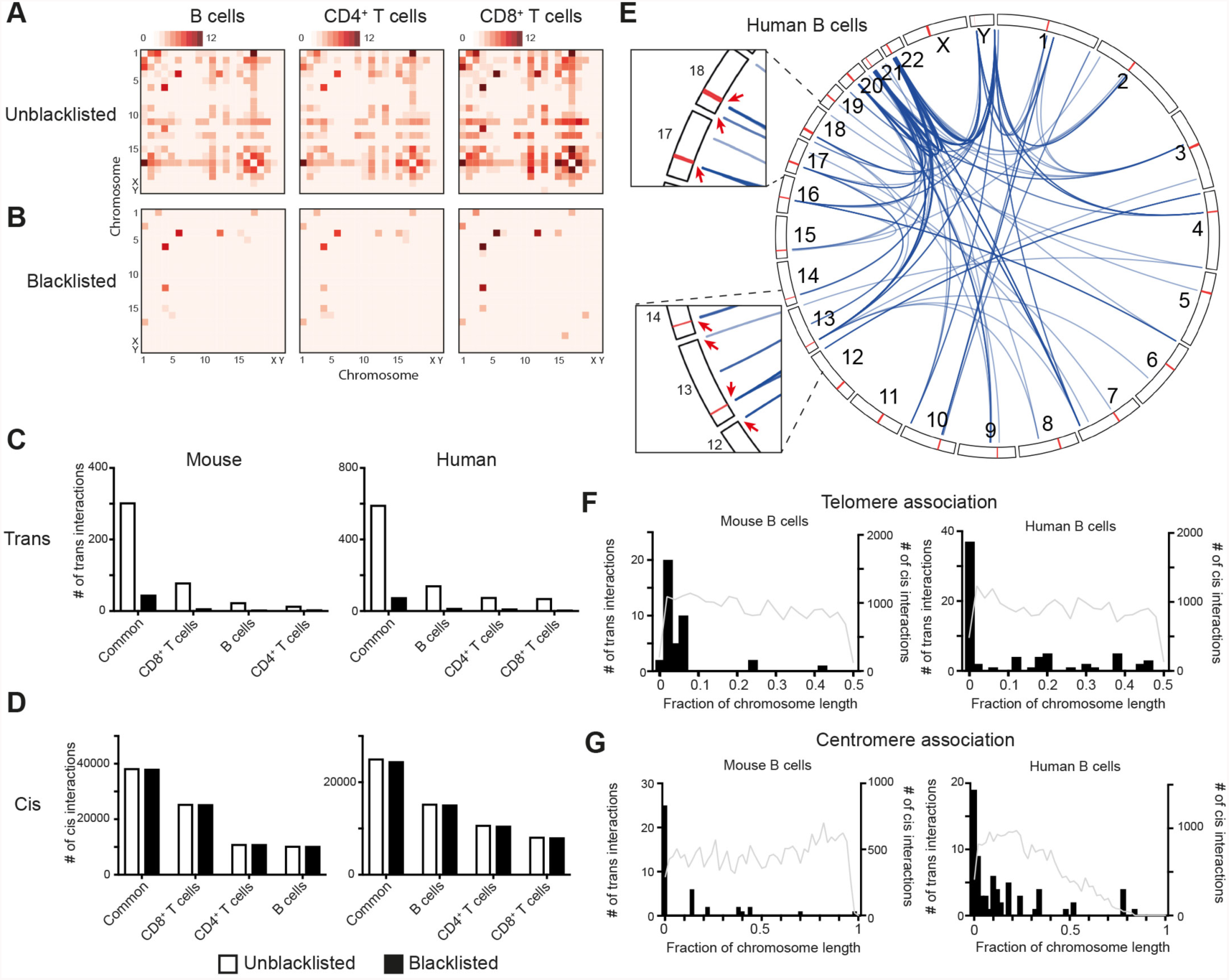
Identification of transchromosomal interactions in mammalian immune cells. **(A)** Heatmap of chromosomes involved in detected transchromosomal interactions in mouse B cells, CD4^+^ and CD8^+^ T cells. **(B)** Heatmap of chromosomes involved in transchromosomal interactions in mouse B cells, CD4^+^ and CD8^+^ T cells after exclusion of interactions associated with blacklisted regions. **(C)** Numbers of transchromosomal interactions common, or unique to murine B cells, CD4^+^ or CD8^+^ T cells before (white) and after (black) exclusion of interactions associated with blacklisted regions. **(D)** Numbers of intrachromosomal interactions common, or unique to murine B cells, CD4^+^ or CD8^+^ T cells before and after exclusion of interactions associated with blacklisted regions. **(E)** Circos plot of transchromosomal interactions in human B cells. Insets show examples of interactions associated with centromeres and telomeres. Centromeres are shown in red. **(F)** Association of mouse or human B cell specific transchromosomal (black histogram) and intrachromosomal interactions (grey line) with telomeres. **(G)** Association of mouse or human B cell specific transchromosomal (black histogram) and intrachromosomal interactions (grey line) with centromeres.

To determine if any of the detected transchromosomal interactions, whether associated with telomeres or centromeres or not, have a gene regulatory function, we examined the relationship between lineage-specific transchromsomal interactions (those found in only one of the cell populations) (Supplemental Table 2) and expression of gene associated with these interactions (*20).* In the mouse, we found that the 15 lineage-specific transchromosomal interactions (3 B cell, 8 CD8+ T cell and 4 CD4+ T cell) overlap only 3 genes *(Cct4, Lars2, Hjurp)* expressed (>5 RPKM) in any of the three lineages and none of these was expressed specifically, or differentially, in the lineage exhibiting the lineage-specific transchromosomal interaction. Similarly, in humans, we found that none of the 38 lineage-specific transchromosomal interactions (18 B cell, 5 CD8+ T cell and 15 CD4+ T cell)(Supplemental Table 2) associated with any protein-coding genes differentially expressed (>5 RPKM) in the lineage exhibiting the lineage-specific transchromosomal interaction. This suggests that none of the detected lineage-specific transchromosomal interactions perform a gene regulatory function in mouse or human B or T cells.

It has been suggested that if transchromosomal interactions were functionally important they would be evolutionarily conserved (2). Therefore, we examined the handful of genes and genomic regions associated with all transchromosomal interactions in mouse and human B and T cells. We found that none of the lineage-specific transchromosomal interactions link orthologous regions in mouse and human.

As we were able to detect transchromosomal interactions, but none of a gene regulatory nature, we examined regions previously reported to be involved in regulatory interactions between chromosomes. We examined our CD4^+^ T cell data for interactions between the previously mentioned LCR region on mouse chromosome 11 and loci encoding the cytokine interferon gamma (Ifng) on chromosome 10 (*11*) and interleukin 17 (IL-17) on chromosome 1 (12). Curiously, no interactions were detected between the LCR and *Ifng* or *IL17* loci in mouse CD4^+^ T cells (Figure 2 A-D). Intrachromosomal interactions at the loci exhibited three-dimensional structure as expected (Figure 2 E-F), indicating that the *in situ* HiC data was of sufficient quality. Similarly, in human CD4^+^ T cells we found no interactions between the LCR and *Ifng* or *IL17 l*oci (Figure 2 G-J). Again, intrachromosomal interactions at the loci were as expected (Figure 2 K-L). These analyses were repeated with raw data (no artefact removal step during data processing) to ensure that reads potentially indicating interactions had not been filtered out. No interactions between the LCR and *Ifng* or *IL17* loci in either mouse or human were detected in the raw, unfiltered data (Supplemental Figure 2 A-D).

**Figure 2:**
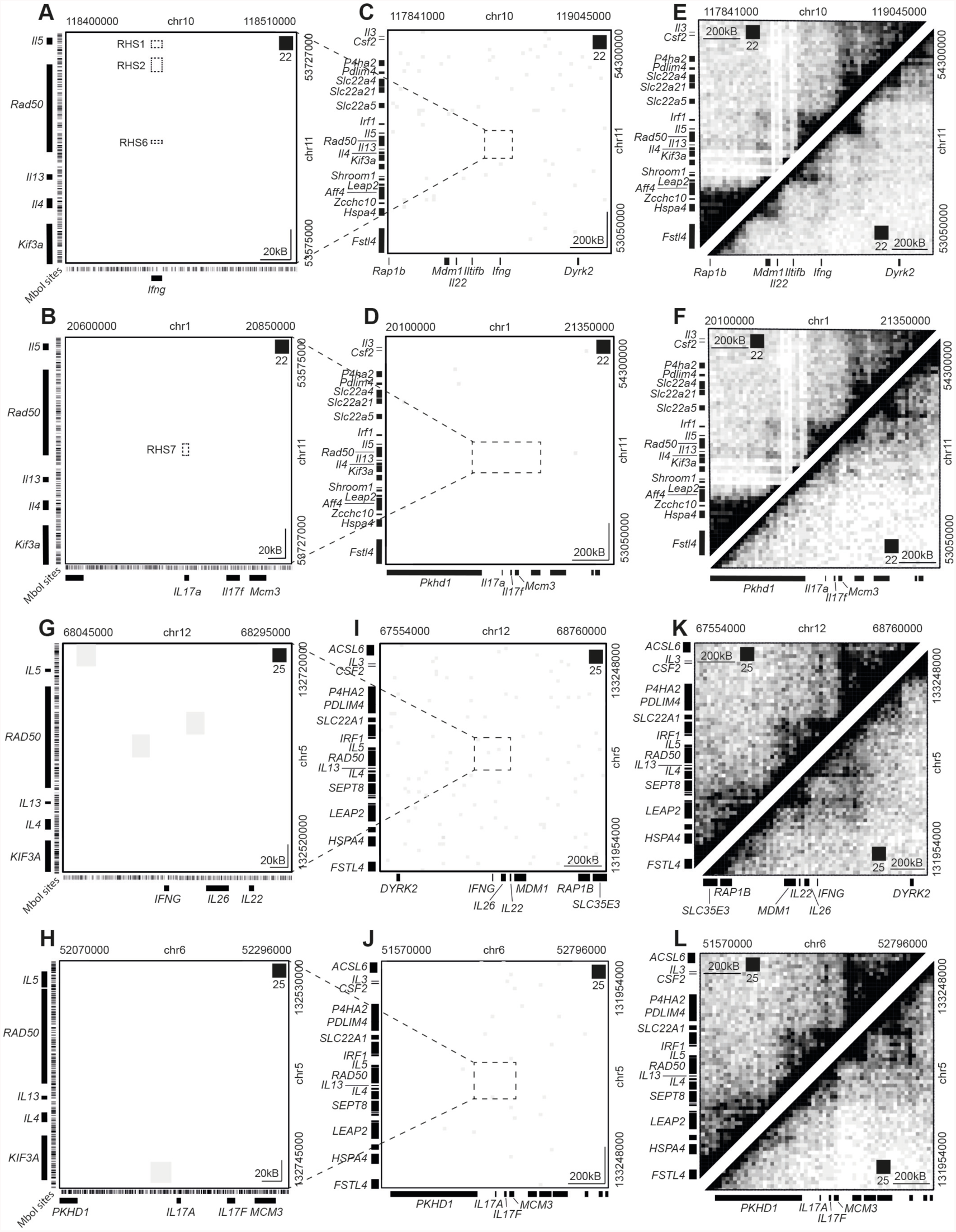
Reported transchromosomal interactions are not detected by *in situ* HiC. **(A)** HiC contact matrix of regions on chromosome 10 and 11 in mouse CD4^+^ T cells previously reported to interact. Dotted squares show regions reported to interact. Colour intensity represents interaction with white being absence of detected interaction and black being intense interaction. **(B)** HiC contact matrix of regions on chromosome 1 and 11 in mouse CD4+ T cells previously reported to interact. Dotted square shows regions reported to interact. **(C)** Expanded HiC contact matrix of regions on chromosome 16 and 11 in mouse CD4+ T cells previously reported to interact. Dotted square encloses the region shown in Figure 2 A. **(D)** Expanded HiC contact matrix of regions on chromosome 16 and 11 in mouse CD4+ T cells previously reported to interact. Dotted square encloses the region shown in Figure 2 B. **(E)** HiC contact matrices showing the detected intrachromosomal interactions in mouse CD4^+^ T cells in the two regions on chromosome 10 and 11 reported to interact in *trans.* **(F)** HiC contact matrices showing the detected intrachromosomal interactions in mouse CD4+ T cells in the two regions on chromosome 1 and 11 reported to interact in *trans.* **(G)** HiC contact matrix of regions on chromosome 12 and 5 in human CD4+ T cells previously reported to interact in mouse CD4+ T cells. **(H)** HiC contact matrix of regions on chromosome 6 and 5 in human CD4+ T cells previously reported to interact in mouse CD4+ T cells. **(I)** Expanded HiC contact matrix of regions on chromosome 12 and 5 in human CD4+ T cells previously reported to interact. Dotted square encloses the region shown in Figure 2 G. **(J)** Expanded HiC contact matrix of regions on chromosome 6 and 5 in human CD4+ T cells previously reported to interact in mouse CD4+ T cells. Dotted square encloses the region shown in Figure 2 H. **(K)** HiC contact matrices showing the detected intrachromosomal interactions in human CD4+ T cells in the two regions on chromosome 12 and 5 reported to interact in *trans.* **(L)** HiC contact matrices showing the detected intrachromosomal interactions in human CD4^+^ T cells in the two regions on chromosome 6 and 5 reported to interact in *trans.*

To determine if the depth of sequencing of our *in situ* HiC had inhibited detection of the previously reported transchromosomal interactions, we examined publically available promoter capture HiC data from human CD4+ T cells (21). The LCR-Ifng or *IL17* interactions were also undetectable in this extremely high-resolution data (Supplemental Figure 2 E, F).

We then attempted to detect another previously reported transchromosomal interaction suggested to occur between the immunoglobulin heavy (*Igh*) and light chain (*Igk*) loci in mouse B cell progenitors (13). Our transchromosomal interaction detection pipeline was applied to *in situ* HiC libraries generated from two B cell progenitors: pro-B cells and immature B cells. Curiously again, using our unbiased, genome-wide approach, we found no interactions between *Igh* on chromosome 12 and *Igk* on chromosome 6 in either B cell progenitor population (Supplemental Figure 2 G, H). Intrachromosomal interactions at both loci were as expected (Supplemental Figure 2 G, H).

In summary, using an unbiased, genome-wide approach we detect neither novel, nor previously reported, gene-regulatory transchromosomal interactions in three dominant mouse and human immune cell populations.

## Discussion

For many years DNA Fluorescent *In situ* Hybridisation (FISH) (*22*) and chromatin conformation capture (3C) (23) were the dominant technologies used to examine chromosomal interactions, whether in *cis* or *trans.* While these technologies have been invaluable in defining the nature of intrachromosomal interactions, their application to interrogate transchromosomal interactions has been less fruitful. Incongruous results from FISH versus 3C within cell types, or in fact from the same technique between studies, has been a persistent issue when examining transchromosomal interactions. For example, as previously mentioned 3C indicated that X chromosomes were in contact prior to X chromosome inactivation (*5–7*) whereas FISH revealed little to no contact (7). In another example, the two studies reporting transchromosomal interactions between Ifg2 and loci on other chromosomes in mouse T cells found no common interactions (*9,10),* while two further studies of the reported interaction in human T cells (8) found no evidence of interaction (*24, 25).*

To address this vexed issue, we used the *in situ* HiC technique to search for transchromosomal interactions across two species and three distinct cell populations. With this unbiased, genome-wide approach, we were unable to detect any conserved, gene regulatory transchromosomal interactions. While our findings are clear and suggest gene regulatory transchromosomal interactions do not function in the mammalian immune system, it is not possible to be totally conclusive about a negative finding. For example, we cannot rule out gene regulatory interactions that are weak, transient, present in highly repetitive regions or in regions without *MboI* restriction sites. Furthermore, because we used only male-derived DNA we could not examine interactions reported to occur between X chromosomes during X chromosome inactivation (26).

Physiologically relevant transchromosomal interactions that are transient and/or weak may not be detectable by *in situ* HiC. However, this does not explain the absence of the interactions between LCR and *Ifng* or *IL17* loci in T cells, or the immunoglobulin loci in B cell progenitors, as these interactions are reported to occur in 40-50% of cells (*11, 13*) and the interactions are reported to be as strong as intrachromosomal interactions (11).

Differences between results presented here and those previously reported are likely due to differences in methodology. Previous studies relied on targeted amplification-dependent chromatin capture techniques and/or DNA FISH. It is increasingly clear that even with the appropriate controls (*23),* a minute amplification bias in a targeting probe combined with the large number of amplification steps required for 3C-based approaches can lead to false positives (2). Furthermore, it has been suggested that up to half of the ligation events in chromatin capture techniques that rely on dilution of the ligation reaction to deduce proximity, such as 3C or ‘dilution’ HiC, link regions of DNA that were not truly associated in the intact nucleus (27). Although DNA-FISH does not exhibit amplification bias, it does suffer from the resolution limitations of light microscopy (250-500nm). Thus it may be that the *Igh* and *Igk* loci in B cell progenitors, or other FISH-demonstrated interactions, frequently lie within hundreds of nanometres of each other, but are nevertheless not sufficiently proximate to be regulatory or chemically crosslinked and thus detected by *in situ* HiC.

While physiologically relevant transchromosomal interactions are observed extensively in fungi (*28*) and *Drosophila* (29), our data suggest that such interactions do not exist in human or murine immune cells. One possible explanation for this difference is that over time the increased frequency of translocation at points of consistent DNA-DNA contact (30–32) induce ultimately advantageous chromosomal rearrangement leaving relevant loci on the same chromosome and thus regulation occurring in *cis.*

In summary, the unbiased, genome-wide *in situ* HiC approach found no evidence for the existence of conserved, lineage-specific, gene regulatory transchromosomal interactions in mammalian immune cells, bringing into question the existence of stable, gene-regulatory transchromosomal interactions underlying immune cell identity.

## Acknowledgements

This work was supported by grants and fellowships from the National Health and Medical Research Council of Australia (G.S. #1058892, L. H. and N.B. #1037321, #1129033, #1080887, A.L. and G.S. #1054618, R.A. and T.J. #1049307, #1100451, T.J. #1124081) and the Australian Research Council (R.A. #130100541). This study was made possible through Victorian State Government Operational Infrastructure Support and Australian Government NHMRC Independent Research Institute Infrastructure Support scheme. The data reported in this paper are tabulated in the Supplemental Materials and archived on the GEO database under accession numbers GSE105776 and GSE105918, GSE9163 for human and mouse respectively. The authors declare no conflicts of interest.

## Supplemental Materials

### Materials and Methods

#### Cell isolation

All animal experiments were performed using C57B/6 male mice at age 6-8 w. Mice were maintained at The Walter and Eliza Hall Institute Animal Facility under specific pathogen-free conditions. Males were randomly chosen from the relevant pool. Animal experiments were performed under the Australian code for the care and use of animals for scientific purposes. Approval for sourcing of human material and experimentation was obtained from The Walter and Eliza Hall Institute’s human research ethics committee (HREC No. 88.03). Results were analysed without blinding of grouping.

Murine CD4^+^ T cells (TCRß^+^ CD4^+^ CD8־ CD44־ CD62L^+^), CD8^+^ T cells (TCRß^+^ CD4־ CD8^+^ CD44־ CD62L^+^), immature B cells (TCRß־CD19^+^ B220^+^ IgM^+^ IgD־) and B cells (TCRß־ CD19^+^ B220^+^ IgM^+^ IgD^+^) were obtained from mechanically homogenized spleens. Pro-B cells were expanded from B220^+^ cells from bone marrow on an OP9 cell layer for 7 days in MEM^+^Glutamax (Gibco) supplemented with 10mM HEPES, 1mM Sodium Pyruvate, 1x non essential amino acids (Sigma) and 50μM β-mercaptoethanol (Sigma). At day 7 the IgM-fraction was isolated using immunomagnetic depletion, following manufacturer’s instructions.

Cryopreserved human peripheral blood mononuclear cells were thawed and stained with antibodies against human αβ TCR, CD4, CD45RA, CD25, CD14, CD16, HLA-DR, and CD19. CD4^+^ T cells (CD14^−^ CD16^−^ TCRαβ^+^ CD4^+^ CD45RA^+^ CD25^−^), CD8^+^ T cells (CD14^−^ CD16^−^ TCRαβ^+^ CD4־ CD45RA^+^ CD25^−^), and B cells (TCRαβ HLA-DR^+^ CD19^+^) and isolated by flow cytometric sorting.

Flow cytometric analyses were performed on BD FACSCanto with sorting on the BD Aria or Influx (BD Bioscience). Antibodies were purchased from BD Bioscience or eBioscience (Supplemental Table 3).

#### HiC

HiC was performed as previously published (*15*). Primary immune cell libraries for both human and mouse were generated in biological duplicate. Libraries were sequenced on an Illumina NextSeq 500 to produce 75bp paired-end reads. Between 160 million and 375 million valid read pairs were generated per sample (Supplemental Table 4). Hi-C sequencing data for mouse pro-B cells and immature B cells was obtained from gene expression omnibus accession number GSE99163.

#### Total RNA isolation

RNA was isolated using the miRNeasy Micro Kit (QIAGEN) following manufacturer’s instructions.

#### RNA-seq analysis

All samples were acquired from two male human donors. Each donor provided one sample per biological condition, giving each condition two replicates. RNA libraries were prepared using an Illumina's TruSeq Total Stranded RNA kit with Ribo-zero Gold (Illumina) according to the manufacturer’s instructions. The rRNA-depleted RNA was purified, and reverse transcribed using SuperScript II reverse transcriptase (Invitrogen). Total RNA-Seq libraries were sequenced on the Illumina NextSeq 500 generating 80 base pair paired end reads. The reads were aligned to the human genome (GRCh38/hg38) using the Rsubread aligner (*33).* The number of fragments overlapping Ensembl genes were summarized using featureCounts (34).

Differential expression analyses were undertaken using the edgeR (35) and limma (36) software packages. Any gene which did not achieve a count per million mapped reads (CPM) of 0.1 in at least 2 samples was deemed to be unexpressed and subsequently filtered from the analysis. Compositional differences between libraries were normalized using the trimmed mean of log expression ratios (TMM) (*37*) method. Counts were transformed to log_2_-CPM with associated precision weights using voom (38). Differential expression was assessed using linear models and robust empirical Bayes moderated t-statistics (39). P-values were adjusted to control the false discovery rate (FDR) below 5% using the Benjamini and Hochberg method. To increase precision, the linear model incorporated a correction for a donor batch effect.

#### HiC data processing

##### Read processing and alignment

Reads from each sample were aligned using the presplit_map.py script in the *diffHic* package v1.4.0 (14). Briefly, reads were split into 5' and 3' segments if they contained the *MboI* ligation signature (GATCGATC), using cutadapt v0.9.5 (*40*) with default parameters. Segments and unsplit reads were aligned to the GRCm38/mm10 build of the *Mus musculus* genome or the GRCh38/hg38 build of the *Homo sapiens* genome using bowtie2 v2.2.5 (41) in single-end mode. All alignments from a single library were pooled together and the resulting BAM file was sorted by read name. The FixMateInformation command from the Picard suite v1.117 (https://broadinstitute.github.io/picard/) was applied to synchronise mate information for each read pair. Alignments were resorted by position and potential duplicates were marked using the MarkDuplicates command, prior to a final resorting by name. This was repeated for each library generated from each sample in the data set. Each BAM file was further processed to identify the *MboI* restriction fragment that each read was aligned to. This was performed using the preparePairs function in *diffHic*, after discarding reads marked as duplicates and those with mapping quality scores below 10. Thresholds were applied to remove artefacts in the libraries, (Supplemental Table 4). Read pairs were ignored if one read was unmapped or discarded, or if both reads were assigned to the same fragment in the same orientation. Pairs of inward-facing reads or outward-facing reads on the same chromosome separated by less than a certain distance (min.inward and min.outward respectively) were also treated as dangling ends and were removed. For each read pair, the fragment size was calculated based on the distance of each read to the end of its restriction fragment. Read pairs with fragment sizes above ~1200 bp (max.frag) were considered to be products of off-site digestion and removed. In this manner, approximately 70-75% of read pairs were successfully assigned to restriction fragments in each library. An estimate of alignment error was obtained by comparing the mapping location of the 3' segment of each chimeric read with that of the 5' segment of its mate. If the two segments were not inward-facing and separated by less than ~1200 bp (chim.dist), then a mapping error was considered to be present. Of all the chimeric read pairs for which this evaluation could be performed, around 1-5% were estimated to have errors, indicating that alignment was generally successful. Technical replicates of the same library from multiple sequence runs were then merged with the mergePairs function of *diffHic.*

##### Data correction and detecting loop interactions

Loop interactions were detected using methods in the *diffHic* package. Read pairs were counted into 50 kbp bin pairs (with bin boundaries rounded up to the nearest *MboI* restriction site) using the *squareCounts* function. Only read pairs mapped to a placed scaffold were included therefore unlocalized and unplaced scaffolds were not included. Mitochondrion read pairs were also excluded. Using the *diffHic* function *correctContact,* we applied an iterative correction procedure (*42*) with some modifications to account for differences in sequencibility, mappability and restriction site frequency in the libraries. The modification to the iterative correction procedure involves performing an additional correction to a bin pairs by the negative binomial mean of the replicate libraries.

Looping interactions were detected using a method similar to that described previously (*15).* Specifically, read pairs were counted in bin pairs for all libraries of a given cell type or condition. For each bin pair, the log-fold change over the average abundance of each of several neighbouring regions was computed. Neighbouring regions in the interaction space included a square quadrant of sides 'x+1' that was closest to the diagonal and contained the target bin pair in its corner; a horizontal stripe of length '2x+1' centred on the target bin pair; a vertical stripe of '2x+1', similarly centred; and a square of sides '2x+1', also containing the target bin pair in the centre. The enrichment value for each bin pair was defined as the minimum of these log-fold changes, i.e., the bin pair had to have intensities higher than all neighbouring regions to obtain a large enrichment value. These enrichment values were calculated using the *enrichedPairs* function in *diffHic,* with 'x' set to 5 bin sizes (i.e., 250 kbp). Putative loops were then defined as those with enrichment values above 0.5, with average count across libraries greater than 10, and that were more than 1 bin size away from the diagonal.

##### Blacklisted regions and removal of centromere and telomere loops

Blacklisted genomic regions were obtained from ENCODE for hg38 and mm10 (*18).* Loops that that had at least one anchor in a blacklisted genomic region were removed. Additionally, loops found with an anchor found within a centromere or telomere region as defined by UCSC genome annotation were removed.

##### Finding overlaps between bin pairs

Overlaps between bin pairs were performed using the overlapsAny function in the InteractionSet package with type = equal and maxgap = 100kb (*43).* This considers an overlap to be present if anchors have a separation of less than the maxgap value and if both anchors of the bins pairs overlap.

#### Promoter capture Hi-C data processing

Promoter capture Hi-C sequencing data for human naive CD4^+^ T cells was obtained from EGA (https://www.ebi.ac.uk/ega) accession number EGAS00001001911. The read processing and alignment was with the same methods as the Hi-C data except, as the restriction enzyme *HindIII* was used in the assay, the reads were split with a ligation signature of AAGCTAGCTT.

#### Visualization of results

Plaid plots were constructed using the plotPlaid function from the *diffHic* package. The range of colour intensities in each plot was scaled according to the library size of the sample, to facilitate comparisons between plots from different samples. Heatmaps of the loops between chromosomes where generated using the R package *gplots* with the function *heatmap.2.* Circos plots were generated with the R package RCircos (44).

**Supplemental Figure 1:**
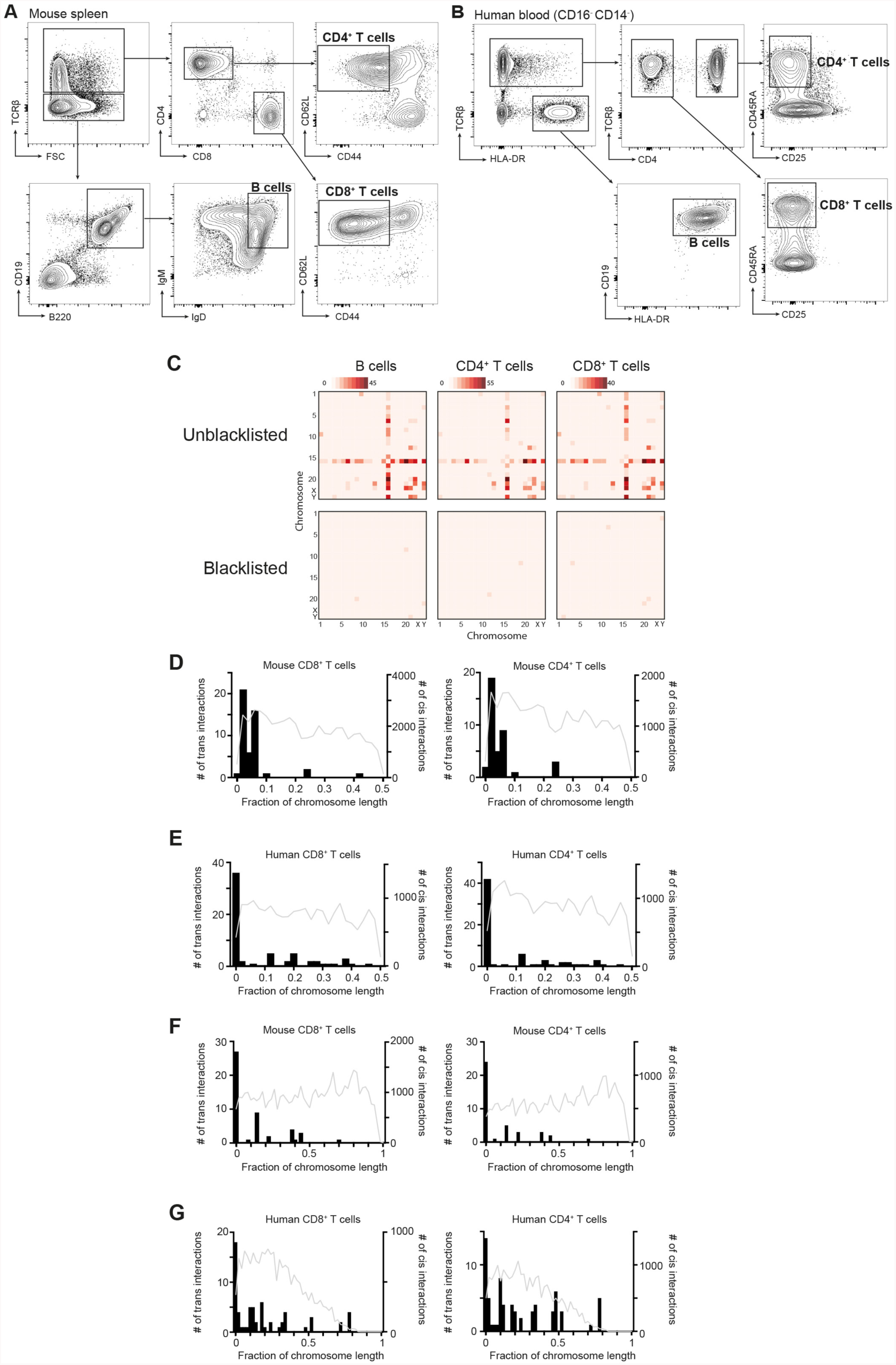
Examination of transchromosomal interactions in mouse and human immune cells. **(A)** Flow cytometry of homogenised C57BL/6 Pep^3b^ mouse spleen stained with antibodies against TCRb, CD4, CD8, CD62L, CD44, CD19, B220, IgD and IgM. CD4^+^ T cells were isolated as TCRb^+^ CD4^+^ CD8־ CD62L^+^ CD44־. CD8^+^ T cells were isolated as TCRb^+^ CD4־ CD8^+^ CD62L^+^ CD44־. B cells were isolated as TCRb־ CD19^+^ B220^+^ IgM^+^ IgD^+^. **(B)** Flow cytometry of human peripheral blood stained with antibodies against TCRb, HLA-DR, CD4, CD45RA, CD25, and CD19. CD4^+^ T cells were isolated as TCRb^+^ CD4^+^ CD45RA־ CD25^+^. CD8^+^ T cells were isolated as TCRb^+^ CD4־ CD45RA־ CD25^+^. B cells were isolated as TCRb־ HLA-DR^+^ CD19^+^. **(C)** Heatmap of chromosomes involved in detected transchromosomal interactions in human B cells, CD4^+^ and CD8^+^ T cells. **(D)** Association of transchromosomal (black histogram) and intrachromosomal interactions (grey line) in mouse CD8^+^ or CD4^+^ T cells with telomeres. **(E)** Association of transchromosomal (black histogram) and intrachromosomal interactions (grey line) in human CD8^+^ or CD4^+^ T cells with telomeres. **(F)** Association of transchromosomal (black histogram) and intrachromosomal interactions (grey line) in mouse CD8^+^ or CD4^+^ T cells with centromeres. **(G)** Association of transchromosomal (black histogram) and intrachromosomal interactions (grey line) in human CD8^+^ or CD4^+^ T cells with centromeres.

**Supplemental Figure 2:**
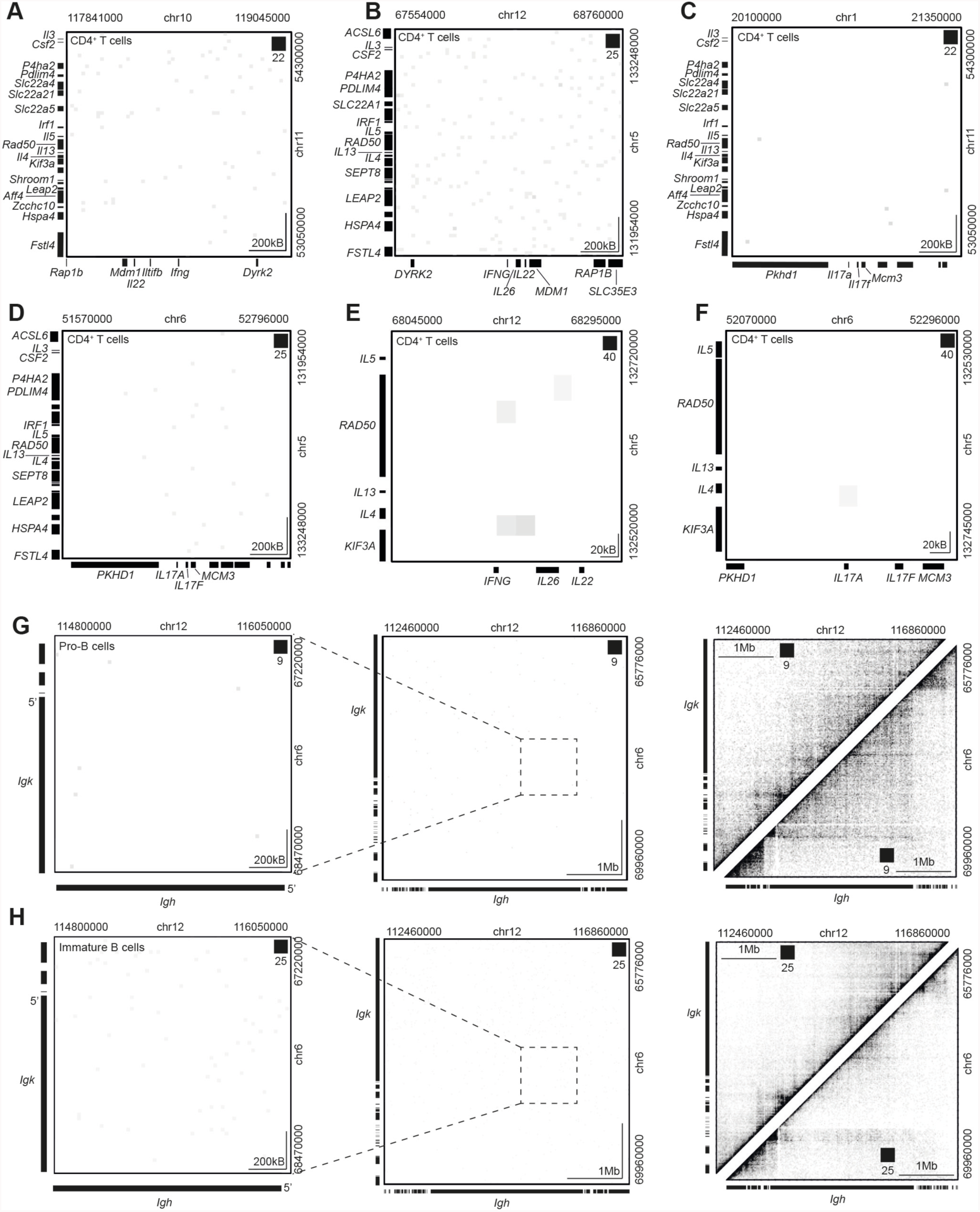
Reported transchromosomal interactions are not detected by in situ HiC, unfiltered in situ HiC or promoter capture HiC. **(A)** HiC contact matrix of unfiltered data of regions on chromosome 10 and 11 in mouse CD4^+^ T cells previously reported to interact. Colour intensity represents interaction with white being absence of detected interaction and black being intense interaction. **(B)** HiC contact matrix of unfiltered data of regions on chromosome 12 and 5 in human CD4^+^ T cells previously reported to interact. **(C)** HiC contact matrix of unfiltered data of regions on chromosome 1 and 11 in mouse CD4^+^ T cells previously reported to interact. **(D)** HiC contact matrix of unfiltered data of regions on chromosome 6 and 5 in human CD4^+^ T cells previously reported to interact. **(E)** Promoter capture HiC contact matrix (21) of regions on chromosome 12 and 5 in human CD4^+^ T cells previously reported to interact. **(F)** Promoter capture HiC contact matrix (21) of regions on chromosome 6 and 5 in human CD4^+^ T cells previously reported to interact. **(G)** HiC contact matrices of regions on chromosome 12 and 6 in mouse pro-B cells previously reported to interact in these cells. The left panel is an expanded plot of the region enclosed by the dotted square in the central panel. The right panel shows the intrachromosomal interactions in the same regions. (H) HiC contact matrices of regions on chromosome 12 and 6 in mouse immature B cells previously reported to interact in these cells. The left panel is an expanded plot of the region enclosed by the dotted square in the central panel. The right panel shows the intrachromosomal interactions in the same regions.

## References and Notes

1. M. Yu, B. Ren, The Three-Dimensional Organization of Mammalian Genomes. 2017, (2017).

2. A. Williams, C. G. Spilianakis, R. A. Flavell, Interchromosomal association and gene regulation in trans. Trends Genet 26, 188–197 (2010).

3. E. J. Clowney et al., Nuclear aggregation of olfactory receptor genes governs their monogenic expression. Cell 151, 724–737 (2012).

4. S. Lomvardas et al., Interchromosomal interactions and olfactory receptor choice. Cell 126, 403–413 (2006).

5. C. P. Bacher et al., Transient colocalization of X-inactivation centres accompanies the initiation of X inactivation. Nat Cell Biol 8, 293–299 (2006).

6. N. Xu, C. L. Tsai, J. T. Lee, Transient homologous chromosome pairing marks the onset of X inactivation. Science 311, 1149–1152 (2006).

7. L. F. Zhang, K. D. Huynh, J. T. Lee, Perinucleolar targeting of the inactive X during S phase: evidence for a role in the maintenance of silencing. Cell 129, 693–706 (2007).

8. J. M. LaSalle, M. Lalande, Homologous association of oppositely imprinted chromosomal domains. Science 272, 725–728 (1996).

9. J. Q. Ling et al., CTCF mediates interchromosomal colocalization between Igf2/H19 and Wsb1/Nf1. Science 312, 269–272 (2006).

10. Z. Zhao et al., Circular chromosome conformation capture (4C) uncovers extensive networks of epigenetically regulated intra- and interchromosomal interactions. Nat Genet 38, 1341–1347 (2006).

11. C. G. Spilianakis, M. D. Lalioti, T. Town, G. R. Lee, R. A. Flavell, Interchromosomal associations between alternatively expressed loci. Nature 435, 637–645 (2005).

12. L. K. Kim et al., Oct-1 regulates IL-17 expression by directing interchromosomal associations in conjunction with CTCF in T cells. Mol Cell 54, 56–66 (2014).

13. S. L. Hewitt et al., Association between the Igk and Igh immunoglobulin loci mediated by the 3' Igk enhancer induces 'decontraction' of the Igh locus in pre-B cells. Nat Immunol 9, 396–404 (2008).

14. Lun A. T., Smyth G. K., diffHic: a Bioconductor package to detect differential genomic interactions in Hi-C data. BMC Bioinformatics 16, 258 (2015).

15. S. S. Rao et al., A 3D map of the human genome at kilobase resolution reveals principles of chromatin looping. Cell 159, 1665–1680 (2014).

16. E. Lieberman-Aiden et al., Comprehensive mapping of long-range interactions reveals folding principles of the human genome. Science 326, 289–293 (2009).

17. A. Chakraborty, F. Ay, Identification of copy number variations and translocations in cancer cells from Hi-C data, bioRxiv, 2017, (2017).

18. E. P. Consortium, An integrated encyclopedia of DNA elements in the human genome. Nature 489, 57–74 (2012).

19. C. Weierich et al., Three-dimensional arrangements of centromeres and telomeres in nuclei of human and murine lymphocytes. Chromosome Res 11, 485–502 (2003).

20. T. S. Heng, M. W. Painter, C. Immunological Genome Project, The Immunological Genome Project: networks of gene expression in immune cells. Nat Immunol 9, 1091–1094 (2008).

21. B. M. Javierre et al., Lineage-Specific Genome Architecture Links Enhancers and Non-coding Disease Variants to Target Gene Promoters. Cell 167, 1369–1384 e1319 (2016).

22. G. T. Rudkin, B. D. Stollar, High resolution detection of DNA-RNA hybrids in situ by indirect immunofluorescence. Nature 265, 472–473 (1977).

23. J. Dekker, K. Rippe, M. Dekker, N. Kleckner, Capturing chromosome conformation. Science 295, 1306–1311 (2002).

24. M. Nogami et al., Relative locations of the centromere and imprinted SNRPN gene within chromosome 15 territories during the cell cycle in HL60 cells. J Cell Sci 113 (Pt 12), 2157–2165 (2000).

25. K. Teller, I. Solovei, K. Buiting, B. Horsthemke, T. Cremer, Maintenance of imprinting and nuclear architecture in cycling cells. Proc Natl Acad Sci U S A 104, 14970–14975 (2007).

26. S. Augui, E. P. Nora, E. Heard, Regulation of X-chromosome inactivation by the X-inactivation centre. Nat Rev Genet 12, 429–442 (2011).

27. T. Nagano et al., Comparison of Hi-C results using in-solution versus in-nucleus ligation. Genome Biol 16, 175 (2015).

28. E. U. Selker, Repeat-induced gene silencing in fungi. Adv Genet 46, 439–450 (2002).

29. I. W. Duncan, Transvection effects in Drosophila. Annu Rev Genet 36, 521–556 (2002).

30. M. R. Branco, A. Pombo, Intermingling of chromosome territories in interphase suggests role in translocations and transcription-dependent associations. PLoS Biol 4, e138 (2006).

31. M. N. Nikiforova et al., Proximity of chromosomal loci that participate in radiation-induced rearrangements in human cells. Science 290, 138–141 (2000).

32. J. J. Roix, P. G. McQueen, P. J. Munson, L. A. Parada, T. Misteli, Spatial proximity of translocation-prone gene loci in human lymphomas. Nat Genet 34, 287–291 (2003).

33. Y. Liao, G. K. Smyth, W. Shi, The Subread aligner: fast, accurate and scalable read mapping by seed-and-vote. Nucleic Acids Res 41, e108 (2013).

34. Y. Liao, G. K. Smyth, W. Shi, featureCounts: an efficient general purpose program for assigning sequence reads to genomic features. Bioinformatics 30, 923–930 (2014).

35. D. J. McCarthy, Y. Chen, G. K. Smyth, Differential expression analysis of multifactor RNA-Seq experiments with respect to biological variation. Nucleic Acids Res 40, 4288–4297 (2012).

36. M. E. Ritchie et al., limma powers differential expression analyses for RNA-sequencing and microarray studies. Nucleic Acids Res 43, e47 (2015).

37. M. D. Robinson, A. Oshlack, A scaling normalization method for differential expression analysis of RNA-seq data. Genome Biol 11, R25 (2010).

38. C. W. Law, Y. Chen, W. Shi, G. K. Smyth, voom: Precision weights unlock linear model analysis tools for RNA-seq read counts. Genome Biol 15, R29 (2014).

39. B. Phipson, S. Lee, I. J. Majewski, W. S. Alexander, G. K. Smyth, Robust hyperparameter estimation protects against hypervariable genes and improves power to detect differential expression. Annals of Applied Statistics 10, (2016).

40. M. Martin, Cutadapt removes adapter sequences from high-throughput sequencing reads. EMBnet. journal 17, pp. 10–12 (2011).

41. B. Langmead, S. L. Salzberg, Fast gapped-read alignment with Bowtie 2. Nat Methods 9, 357–359 (2012).

42. M. Imakaev et al., Iterative correction of Hi-C data reveals hallmarks of chromosome organization. Nat Methods 9, 999–1003 (2012).

43. A. T. Lun, G. K. Smyth, csaw: a Bioconductor package for differential binding analysis of ChIP-seq data using sliding windows. Nucleic Acids Res 44, e45 (2016).

44. H. Zhang, P. Meltzer, S. Davis, RCircos: an R package for Circos 2D track plots. BMC Bioinformatics 14, 244 (2013).

